# piRNAs are abundant in the early embryo of the crustacean *Parhyale hawaiensis*

**DOI:** 10.1101/2025.05.30.656998

**Authors:** Llilians Calvo, Tom Pettini, Guillem Ylla, Matthew Ronshaugen, Sam Griffiths-Jones

## Abstract

PIWI-interacting RNAs (piRNAs) are a group of short (∼21-31 nucleotide) non-coding RNAs that protect the germline of metazoans against the activity of genomic parasites known as transposable elements (TEs). Although originally discovered in *Drosophila* where they are germ-line restricted, recent studies in arthropods have shown piRNAs are often expressed also in somatic cells suggesting they might have originally evolved as a somatic immune system protecting against the detrimental action of TEs in the last common ancestor. Beyond their classic TE-silencing roles piRNAs functions range from sex determination in moths, to degradation of maternal mRNAs during the maternal-to-zygotic transition, thus highlighting the need for more sampling across the tree of life, particularly in underrepresented speciose rich subphyla such as the crustaceans were to date only two representatives have been analysed. In this study we sequenced and analysed putative piRNAs in the amphipod *Parhyale hawaiensis*, throughout a time-course of embryogenesis. Initially, during early embryogenesis piRNAs-mapping reads were abundant, maternally loaded piRNAs and showed the hallmark signatures of piRNA processing suggesting active targeting of TEs. Interestingly, this initial high content was followed by a dramatic loss of piRNA-mapping transcripts and signatures during mid-embryogenesis shortly after germ cell specification. This was confirmed by in-situ hybridization of key mRNAs coding for piRNA pathway proteins which revealed their expression becomes restricted specifically to the germ cells by early germband formation, providing an explanation for the observed reduction in piRNA abundance at later stages, and suggesting that piRNAs are absent in *Parhyale* post embryonic somatic cells.

## Introduction

One of the mechanisms that controls the activity of selfish invasive elements in metazoan genomes, particularly in the germline, involves a class of small non-coding RNAs known as PIWI-interacting RNA (piRNAs). piRNAs are typically 21-33 nucleotides (nt) in size, and associate with proteins of the Argonaute family, to silence transposable elements (TEs) either at the transcriptional or the post-transcriptional level^1^.

Much of the fundamental understanding of piRNA biology have come from work in a limited range of model organisms, dominated by *Drosophila*, *Caenhorhabditis elegans*, and mouse. In *Drosophila,* piRNAs are found to originate from two pathways: the primary pathway and the ping-pong cycle (secondary pathway)^2,3^. piRNAs derived from the primary pathway are expressed from genomic regions known as piRNA clusters. piRNA clusters are often located near euchromatin boundaries and function as a defense mechanism: when a transposable element (TE) inserts into a piRNA cluster, it initiates the production of piRNAs that, through sequence complementarity, recognize and silence the TE^4,5^. In brief, cluster-derived piRNAs once transported into the cytoplasm are O-methylated in their 3’ end by a small RNA 2′-O-methyltransferase DmHen1/Pimet^6,7^, which produces mature piRNAs that are loaded onto Piwi protein to produce the Piwi-piRISC complexes. These Piwi-piRNA complexes can then be imported back into the nucleus to guide histone methylation (H3K9me3) by methyltransferases such as Eggless and the heterochromatin protein HP1a. Methylation induces heterochromatin formation thereby silencing target genes^8–10^.

In *Drosophila*, cytoplasmic Aub-bound piRNAs resulting from the primary pathway initiate a second biogenesis process of piRNA production/amplification known as the ping-pong cycle. AGO3 and Aub act in a complementary fashion to slice and silence active transposon mRNAs and at the same time produce more piRNAs. Aub-piRNAs are complementary to TE transcripts and are used as guides to mediate the cleavage of TE transcripts resulting in new piRNA sequences that are recognized by and bind to the AGO3 protein. In turn, AGO3-piRNAs recognize and cleave cluster-derived piRNAs to generate sequences that are very similar to piRNAs from the primary pathway and are therefore recognized by Aub. The Vasa protein promotes the transfer of piRNAs between ping-pong partners^11,12^. piRNAs produced from the ping-pong cycle can be recognized by characteristic properties that result from the processing mechanism: overlapping pairs of piRNAs have a signature 1U/10A bias and a 10nt 5′-5′ overlap as a consequence of Aub and AGO3 slicer activity between nucleotides 10 and 11 of target RNAs^2,3^.

While they are mostly known for their role controlling transposons, piRNAs have also been shown to act on coding genes, mostly via their 3′UTRs^13^. For example, during the maternal to zygotic transition (MZT) in *Drosophila*, when maternal mRNAs are degraded, piRNAs mediate Aub-dependent destabilizes those mRNAs^14^. piRNAs have also been shown to guide translational repression and decay of *nanos* mRNAs, which when impaired, caused aberrant nanos protein expression and resulting segmentation defects^15^.

Studies in different animals have revealed a variety of adaptations in piRNA function across the metazoan tree. However, striking differences have also been found between more closely related species belonging to the same phylum, for instance within the arthropods. Initial studies in *Drosophila* found piRNAs act as a defence mechanism against transposon activity, almost exclusively in the germ-line^16^. In ovaries, the primary pathway is active in both somatic cells and the germ cells, whereas ping-pong amplification only occurs in the germ cells^2,16^. This segregation in the gonads can also be observed in the localization of proteins directing the biogenesis pathway: somatic ovarian cells only express Piwi protein, whereas germ cells express all three proteins involved in the ping-pong pathway: Piwi, Aub and AGO3^17^. However, more recent studies in *Drosophila* have reported the expression of piRNAs in somatic tissues such as the brain^18^ but not in the thorax of flies^19^. In *Apis mellifera* (honeybee) where there is an uncoupling between reproduction and labour, piRNAs have been found to be more abundant in the reproducing castes, and are noticeably higher in the haploid males where the effects of TE insertions would be most deleterious^20^. However, in other Hymenoptera, recent studies have found piRNAs to be absent from the male germ line of the bumblebee *Bombus terrestris*^19^. In the silkworm *Bombyx mori*, piRNAs have been shown to play a decisive role in sex determination. In this species, females are heterogametic and their sex is determined by the presence of two chromosomes (WZ). The W chromosome carries the female-specific piRNA *Feminizer* (*Fem*), which acts as the feminizing factor. *Fem* piRNAs induce cleavage of the mRNA produced from the protein coding gene *Masculinizer* (*Masc*) on the Z chromosome, thereby preventing masculinization of the embryos^21^. Recent studies across several different arthropod species now support the expression of piRNAs outside the germ line. For instance, somatic piRNAs have been described in multiple chelicerates, insects and the myriapod *Strigamia maritima*^19,22–25^. Therefore, it is now considered that somatic piRNA expression is the ancestral state of all arthropods, and that the somatic pathway has been subsequently lost in several species^19^. However, the documented losses of somatic piRNAs are spread throughout the arthropod phylogenetic tree, making predictions regarding somatic piRNA presence in any particular arthropod species difficult. The function of somatic piRNAs remains elusive and controversial in many systems studied to date^26^.

Crustacea is a predominantly aquatic and highly abundant subphylum of the Arthropoda, comprising an estimated 67,000 species^27^ and accounting for around 0.94Gt of the earth biomass^28^. Analysis of crustacean piRNAs has so far been limited to the terrestrial isopod *Armadillium vulgare,* for which no somatic piRNAs were detected^19^. There have been no studies of piRNAs in aquatic crustacea. Therefore, the role of piRNAs in this entire subphylum remains unknown. Here, we use high-throughput sequencing of small RNAs in the crustacean *Parhyale hawaiensis* to investigate the presence of somatic piRNAs during development. We find clear evidence for both primary and secondary piRNAs in the early embryo, suggesting the ping-pong pathway is active at these stages and that the early embryo is pre-loaded with piRNAs. However, we do not find abundant piRNAs in somatic cells at later stages. Using whole transcriptome RNA-Seq data, we show that mRNAs important to the piRNA biogenesis pathway are also abundant in the early embryo. These data suggest that piRNAs are indeed involved in *Parhyale* early embryonic development, but that their role diminishes later in embryogenesis, or are restricted to just a specific subset of cells, possibly germ cells. In support of this idea, we show that expression of *vasa* and *piwi* is restricted to just the germ cells after germband formation in *Parhyale*. This is only the second study examining piRNAs in crustaceans and the first to analyse dynamic expression in a marine crustacean through embryonic development. The two studies together provide no evidence for widespread expression of somatic piRNAs in this subphylum.

## Materials and Methods

### RNA isolation and library preparation

Precisely staged *Parhyale* embryos were collected and stored in RNA later (Sigma-Aldrich). RNA extraction was performed using the SPLIT RNA Extraction Kit (Lexogen) following the manufacturer’s instructions, splitting RNAs into two fractions: short (<150nt) and long (>150nt). Small RNA libraries (4 replicates per time-point) were built using the Small RNA-Seq Library Prep Kit (Lexogen), and libraries were purified using the purification module with magnetic beads from Lexogen. Long RNA-Seq libraries (2 replicates per time-point) were built using the TruSeq Stranded mRNA HT Sample Prep Kit (Illumina). Sequencing for all libraries was performed at the University of Manchester Genomic Technologies Facility.

### Identification of putative piRNA reads

RNA-Seq reads were trimmed using Cutadapt v1.18^29^ and reads ranging from 25 to 35nt in length were retained. These reads were mapped to the *Parhyale* genome (Phaw_5.0-GCA_001587735.2) using bowtie1^30^ allowing 1 mismatch (bowtie -v 1 -S -a –best –strata -a - al). Reads that mapped to the genome were filtered to remove potential tRNA, rRNA and mature microRNA sequences, using tRNAs previously predicted with tRNAscan-SE (v2.0), crustacean rRNA downloaded from RNAcentral (https://rnacentral.org, v17)^31^, and microRNAs predicted previously by us^32^ (bowtie1 alignment, allowing 3 mismatches for tRNAs and rRNAs, and 0 mismatches for microRNAs). The remaining reads were considered putative piRNAs and retained for further analysis.

### piRNA cluster prediction

piRNA clusters were predicted using reads that map with 0 mismatches only once to the genome (bowtie -v 0 -m 1 -a –best –strata) and the software Protrac v2.4.3^33^ with the following criteria: minimum size for a piRNA cluster: 2.5 kb (-clsize 2500), p-value for minimum number of (normalized) read counts per kb: 0.05 (-pdens 0.05), minimum fraction of (normalized) read counts that have 1T (1U) or 10A: 0.5 (-1Tor10A 0.5), minimum fraction of hits with 1T (U) and 10A: 0.33 (-1Tand10A 0.33), fraction of (normalized) read counts that map to the main strand: 0.5 (-clstrand 0.5), minimum number of sequence hit loci per piRNA cluster: 1000 (- clhits 1000), size of the sliding window: 10000 bp (-swsize 10000), increment of the sliding window: 1000 (-swincr 1000).

### Analysis of putative piRNAs locus of origin

Putative piRNAs were mapped to four potential origins in the *Parhyale* genome: Transposable elements, piRNAs clusters, mRNAs and 3’UTRs using (bowtie -v 3 -a -best). A 151 *Parhyale* consensus transposable elements sequences were obtained from Repbase v25.03. piRNA clusters (piClusters) were predicted as showed above whereas *Parhyale* mRNAs and *Parhyale* 3’UTRs were previously annotated by us^34^.

Ping-pong signature were calculated by quantifying 5′-5′ reads overlap from 1nt to 35nt using the script signature.py^35^. Reads with 10nt overlap mapping on opposite strands were considered to be generated by the ping-pong mechanism.

Transposable elements quantification were performed by using previously generated RNA-Seq reads^34^. In brief, RNA-Seq reads where trimmed, mapped to the genome (Phaw_5.0-GCA_001587735.2) and mappers were subsequently mapped to consensus transposable element using Bowtie2 (-N 1 –local). Mapping reads were quantified and normalized using DESeq2 v1.28.1 and FeatureCounts v2.0.1. Additionally, putative piRNA reads mapping to *Parhyale* consensus transposable elements were also quantified and normalized using featureCounts v2.0.1^36^ and DESeq2 v1.28.1^37^. All figures were plotted in RStudio using the ggplot2^38^ and ComplexHeatmap packages^39^.

### smiFISH detection of *vasa* and *piwi* transcripts

*Parhyale* embryo fixation and smiFISH staining were performed as detailed in^40^ *vasa* and *piwi* gene-specific smiFISH probe sequences are provided in Supplementary Table 1. *vasa* probes were labelled with Quasar 570, and *piwi* probes with CalFluor 610. smiFISH images were taken using a Leica TCS SP8 AOBS inverted gSTED microscope with a 40x/1.3 HC PL APO (oil) objective. Image stacks were acquired with 400nm z-interval over a total depth of 16um, with the following settings: format 2048×2048, speed 400Hz unidirectional, sequential line scanning, line averaging 8, frame accumulation 3, pinhole 1 airy unit. Each channel was gated 1.0-6.0. DAPI excitation 405nm, laser 5%, collection 415-480nm, CalFluor 610 excitation 590nm, laser 15%, collection 600-642nm, Quasar 570 excitation 548nm, laser 15%, collection 558-585nm. Z-stacks were stabilized through z to correct any imaging drift, and deconvolved using Huygens Professional v18.04. Maximum projections were generated in FIJI v 2.0.0-rc-49/1.51d, and combined into RGB images in Adobe Photoshop CS6.

### Data availability

All RNA sequencing data and quantifications were deposited in the Gene Expression Omnibus (GEO) database under accession number GSE178877.

## Results

### Identification of putative piRNAs in *Parhyale*

Somatic piRNAs have been described in many species of arthropod, and the current thinking is that the piRNA pathway was active in the soma of the last common ancestor of all arthropods^41^. However, the only crustacean investigated to date, *A. vulgare*, failed to show any somatic piRNAs^19^. Therefore, in order to investigate the variation and evolution of the active piRNA pathway in the soma, we collected and analysed small RNA deep sequencing data across embryonic development in the marine crustacean *Parhyale*. We built 28 small RNA libraries covering seven different time-points spanning the full course of embryonic development from the very early embryo at the 1 to 8 cell stages (S1-4), to late embryos (S24-30) (Figure 1A). The final stage, S30, is the fully formed hatchling, which resembles the adult form (*Parhyale* is a direct developer). In order to profile piRNAs through *Parhyale* development we used a discrimination and elimination process, whereby reads mapping to tRNAs, rRNAs and microRNAs were sequentially removed, to leave retained reads annotated as putative piRNAs (Figure 1B). Sequence logo analysis confirmed that the remaining reads show the expected signatures of piRNAs, specifically the 1U/10A bias (1T/10A for DNA reads) that results from the piRNA biogenesis pathway (Figure 1C).

**Figure 1:**
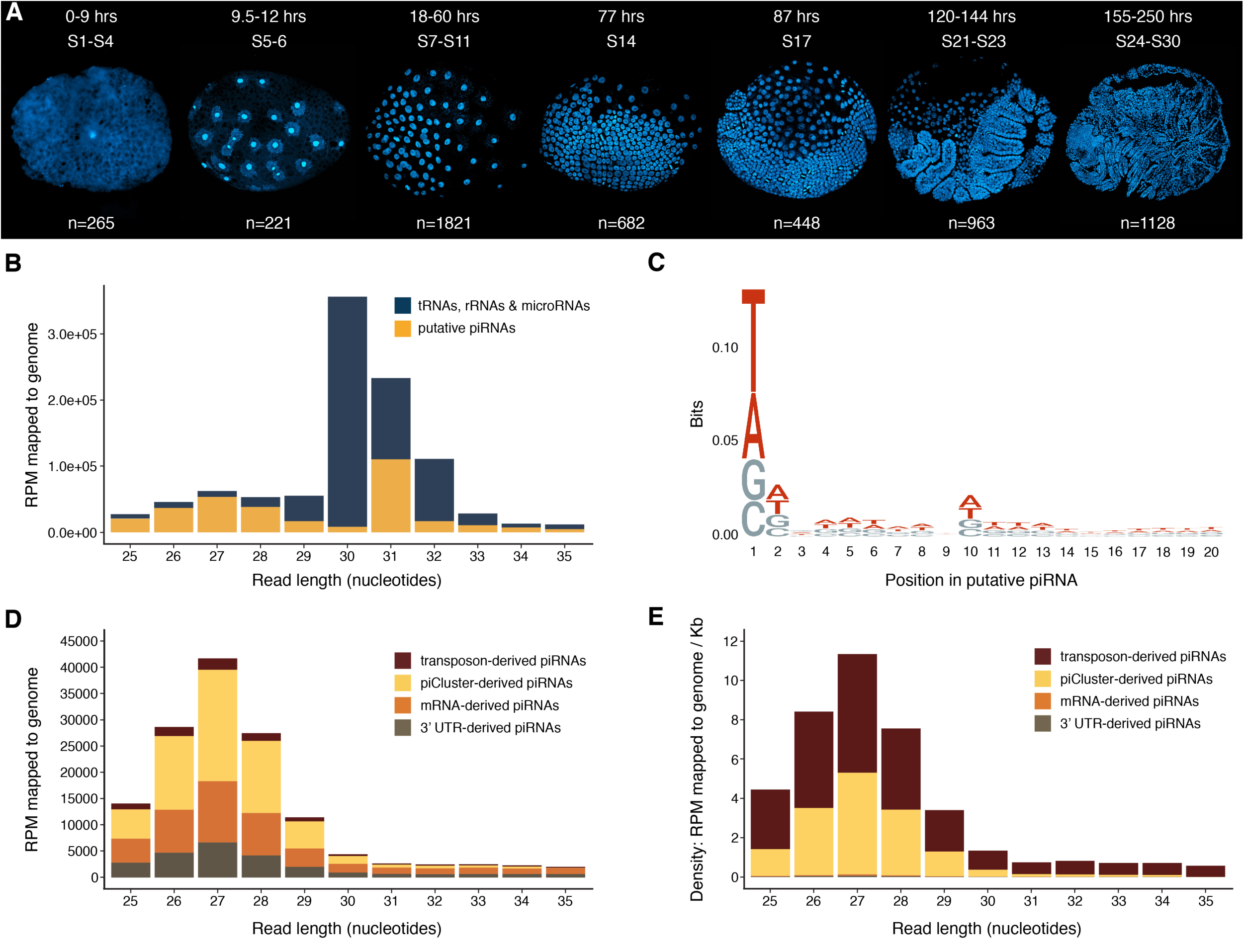
Identification of putative piRNAs and piRNA pathway proteins from *Parhyale* developmental RNA-Seq. (A) DAPI pictures of the seven key developmental time-points used to generate RNA-Seq libraries, spanning the full duration of *Parhyale* embryonic development. Developmental stages, with the number of hours post-fertilization, and the number of embryos pooled for RNA extraction for each time-point are indicated. S1-4 includes 1 to 4 cell stages, and therefore exclusively maternally loaded RNAs. S5-6 marks the maternal-to-zygotic transition, and during S7-S11, embryonic cells migrate and are segregated from the yolk cells. Stages 14 and 17 are both stages of germ band extension, while S21-23 and S24-30 are the stages during which limb development and organogenesis occur. (B) Nucleotide length distribution of 25-35nt reads mapping to the genome, shown as reads per million (RPM) mapped to genome. The proportion of reads of known origin, mapping to transfer RNAs (tRNAs), ribosomal RNAs (rRNAs) or microRNAs are shown in dark blue; remaining reads of unknown origin were considered putative piRNAs (yellow). (C) Sequence logo of the first 20 nucleotides of all putative piRNAs, pooled across all developmental time-points. (D) Nucleotide length distribution of 25-35nt putative piRNAs, shown as reads per million (RPM) mapped to genome. Colour coding indicates the genomic origin. (E) Nucleotide length distribution of 25-35nt putative piRNAs, showing the density of piRNA production as reads per million (RPM) mapped to genome per kb. Colour coding indicates the genomic origin. (F) Heatmap showing expression level in transcripts per million (TPM) of the mRNAs coding for key proteins involved in small RNA pathways, across the seven developmental time-points.

### *Parhyale* piRNAs are largely derived from piRNA clusters and TEs

Previous studies have found that piRNAs are predominantly derived from transposable elements (TEs) and piRNA clusters (piClusters), and also to a lesser degree from mRNAs and 3′UTRs^42,43^. To assess the genomic origin of the putative *Parhyale* piRNAs, we mapped their reads to these four established genomic origins. *Parhyale* consensus TEs were obtained from Repbase, whereas piClusters, mRNAs (CDS) and 3′UTRs were annotated by us. To predict piClusters, putative piRNAs that mapped to a single genomic position were used, and clusters closer than 10kb from one another were merged, resulting in a total of 108 piClusters of primary piRNA production. The majority of predicted piClusters were bidirectionally transcribed, thus producing putative piRNAs from both strands. In *Drosophila*, bidirectional piClusters have been predominantly associated with expression from germ cells, whereas unidirectional piClusters have been observed more often in the somatic cells of the ovaries surrounding the germ cells^44^. Putative piRNA reads mapping to piClusters, Repbase TEs, mRNAs and 3′UTRs are presented in Figure 1D. The large majority of putative piRNA reads mapping to these sites are 25-29nt in length, with a modal size of 27nt. Across sizes 25 to 29nt, the piRNA reads are derived from the four genomic origins in the following order from most to least: piClusters, mRNAs, 3′UTRs, TEs. However, mRNAs and 3′UTRs represent orders of magnitude more sequence than TEs and piClusters (sum lengths: TEs 359.8kb, piClusters 4099kb, 3′UTRs 92299.2kb, mRNAs 180811.4kb). When normalised for these lengths, the density of production of 25 to 29nt sequences from TEs and piClusters is similar, and much higher (∼50 to 120 times) than the density from mRNAs and 3′UTRs (Figure 1E). This high density of piRNA production from TEs and piClusters relative to mRNAs and 3′UTRs is consistent with previous studies^45^.

### piRNAs are abundant during *Parhyale* early embryogenesis

To investigate changes in piRNA prevalence and potential activity across embryogenesis, and to test whether piRNA sequence characteristics vary with their genomic origin, we analysed the size profile, sequence logo and ping-pong signatures of putative piRNAs derived from *Parhyale* consensus TEs, piClusters, mRNAs and 3′UTRs across each of the seven developmental time-points (Figures 2). The plots in Figures 2A show the expression level as reads per million mapped to the genome (RPM) of putative piRNAs ranging from 25-35nt, across each time-point. For all four genomic origins, the highest levels of expression are found in the first three time-points covering the first 60h of embryogenesis. Several events take place during this window, initially only maternally provided transcripts are present (S1-S4), however 9.5-12h after fertilization the embryo becomes transcriptionally active and independent from the maternally deposited material (S5-S6). Finally, in the next 18-60h post fertilization corresponding to the third time-point both gastrulation and germband formation take place (S7-S11). Thereafter, piRNAs levels are dramatically reduced in all future developmental stages. The size profiles of piRNA reads for these first three time-points are highly consistent across the different genomic origins, with a modal length of 27nt in all cases (Figures 2A, time-points S1-4, S5-6 and S7-11). The majority of reads that overlap TEs are antisense to the TE annotation (Figure 2A). This is consistent with ping-pong amplification, indicating that piRNA-mediated silencing of TEs is active in the *Parhyale* early embryo^46^.

**Figure 2:**
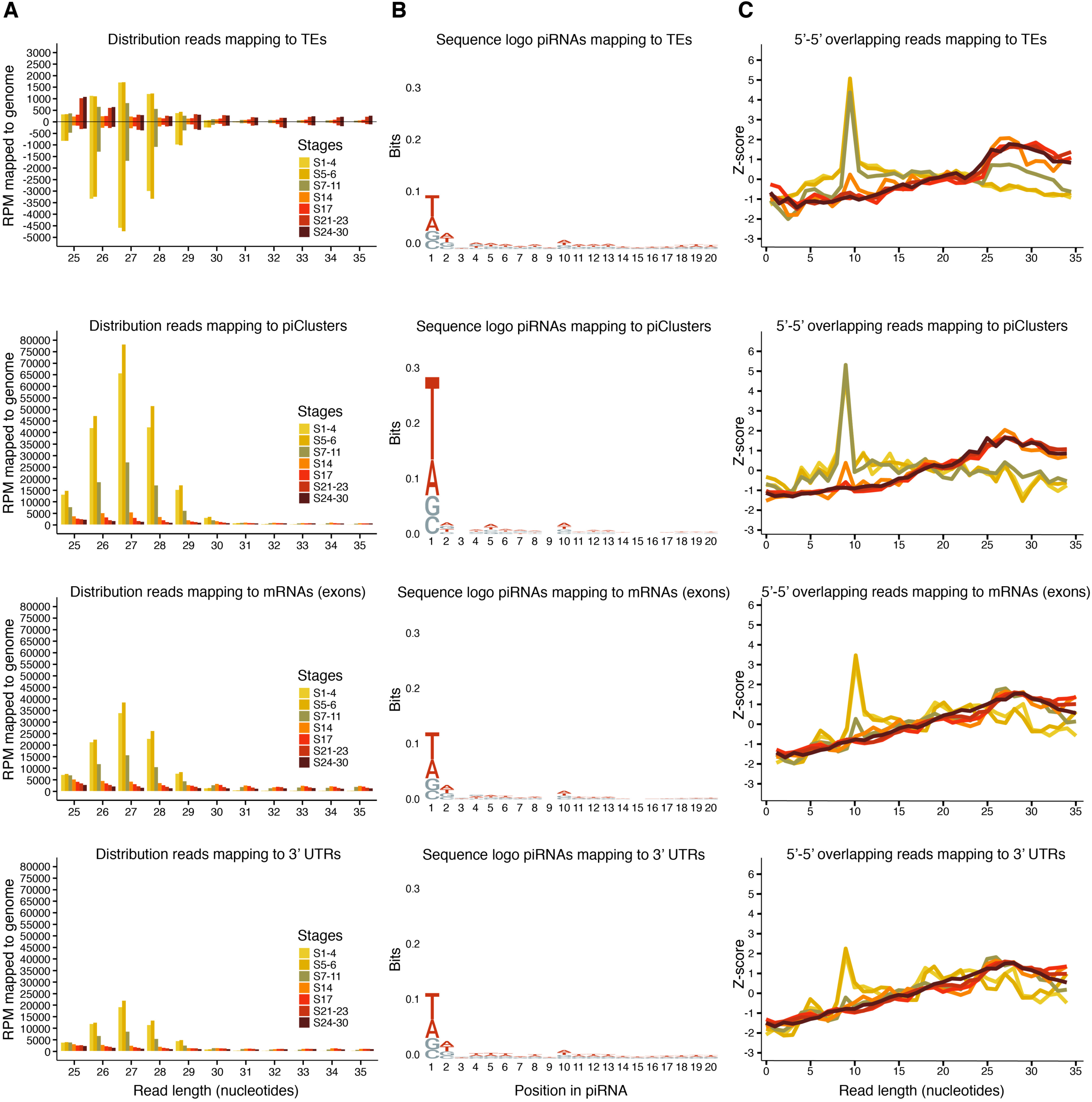
Characterization of piRNAs mapping to consensus transposable elements (TEs), piRNAs custer (piCLusters), mRNAs and 3’ UTRs through embryogenesis. (A) Nucleotide length distribution of 25-35nt putative piRNAs mapping to transposable elements (TEs), piRNAs clusters (piClusters), mRNAs and 3’UTRs shown as reads per million (RPM) mapped to the genome. (B) Sequence logos of the first 20 nucleotides of putative piRNAs. (C) Ping-pong analysis for 25-35nt putative piRNAs mapping sense and antisense to TEs, piClusters, mRNAs and 3’UTRs. Z-scores are plotted for differing degrees of 5′-5′ overlap (1-35nt) between overlapping sense/antisense piRNA pairs.

### piRNAs produced during early embryogenesis from piClusters and TEs show strong signatures of ping-pong amplification

For all four genomic origins, sequence logos reveal that the first three time-points show the 1U/10A enrichment signature typical of piRNAs produced by the ping-pong pathway. This signature deteriorates by S14 and is lost from S17 onwards (Figures 2B). We also analysed all pairs of overlapping sense/antisense reads, and calculated the enrichment of each degree of overlap between 5′ ends (Figures 2C). A peak 5′-5′ overlap at 10nt is indicative of piRNA amplification via the ping-pong pathway^12,41^. Figures 2C show that reads from all four genomic origins display this ping-pong signature, but with temporal variability. Reads derived from piClusters and TEs show strong peaks at 10nt 5′-5′ overlap at the first three time-points, but these peaks are greatly diminished by S14, and lost at all later time-points. For mRNAs and 3′UTRs, this loss of the signature occurs one time-point earlier, with clear peaks only at S1-4 and S5-6, minimal peaks at S7-11, and complete loss of the signature by time-point S14. At the earliest two time-points (S1-4 and S5-6), the 10nt peak is much more pronounced for piRNAs derived from TEs and piClusters (z-scores ∼5) compared with mRNAs and 3′UTRs (z-scores ∼3 and ∼2 respectively). Together, this indicates that piRNA amplification via the ping-pong pathway is more associated with those piRNAs derived from piClusters and TEs, while mRNAs and 3′UTRs likely produce a higher proportion of primary piRNAs.

Together, piRNA total read counts, sequence logos and ping-pong analysis all indicate abundant piRNAs produced via the ping-pong pathway in early embryogenesis, up to S11. By time-point S14, piRNA reads dramatically drop, the high z-score at 10nt 5′-5′ overlap is lost, and sequence logos lose their piRNA signatures, suggesting that piRNAs are only abundant in the early embryo. The presence of the highest read counts and strong ping-pong signatures at the earliest time-point (S1-4) indicates that secondary piRNAs are likely maternally loaded into the egg.

### Maternal TE transcripts are pre-loaded into the egg, and targeted by piRNAs

We examined more closely the developmental expression profiles of individual TEs and the piRNAs that target those TEs. The heatmap presented in Figure 3A shows the expression level across the first three developmental time-points (S1-4, S5-6 and S7-11), of 151 individual *Parhyale* consensus TEs generated from RNA-Seq reads^34^, and of the piRNAs generated from small RNA-Seq that map to the forward and reverse strand of each TE. We found that 96 % of consensus TEs sequences annotated in *Parhyale* were maternally loaded into the early embryo and present even before zygotic transition was activated (S1-4). The expression levels across the TEs are highly variable, yet the levels of each TE transcript generally remain consistent across the three time-points, with only a few TEs displaying increases in expression (indicative of zygotic transcription) or reduction (indicative of TE RNA degradation). For each TE, we identified expression of piRNAs mapping to both sense and antisense strands, with the general trend of higher levels mapping to the antisense strand indicating that TEs are being silenced in the early embryo^47^. Consistent with these findings, as the embryo becomes transcriptionally independent following zygotic genome activation the correlation between the expression of TEs and the piRNAs that target them also increases (Supplemental figure 1.).

**Figure 3:**
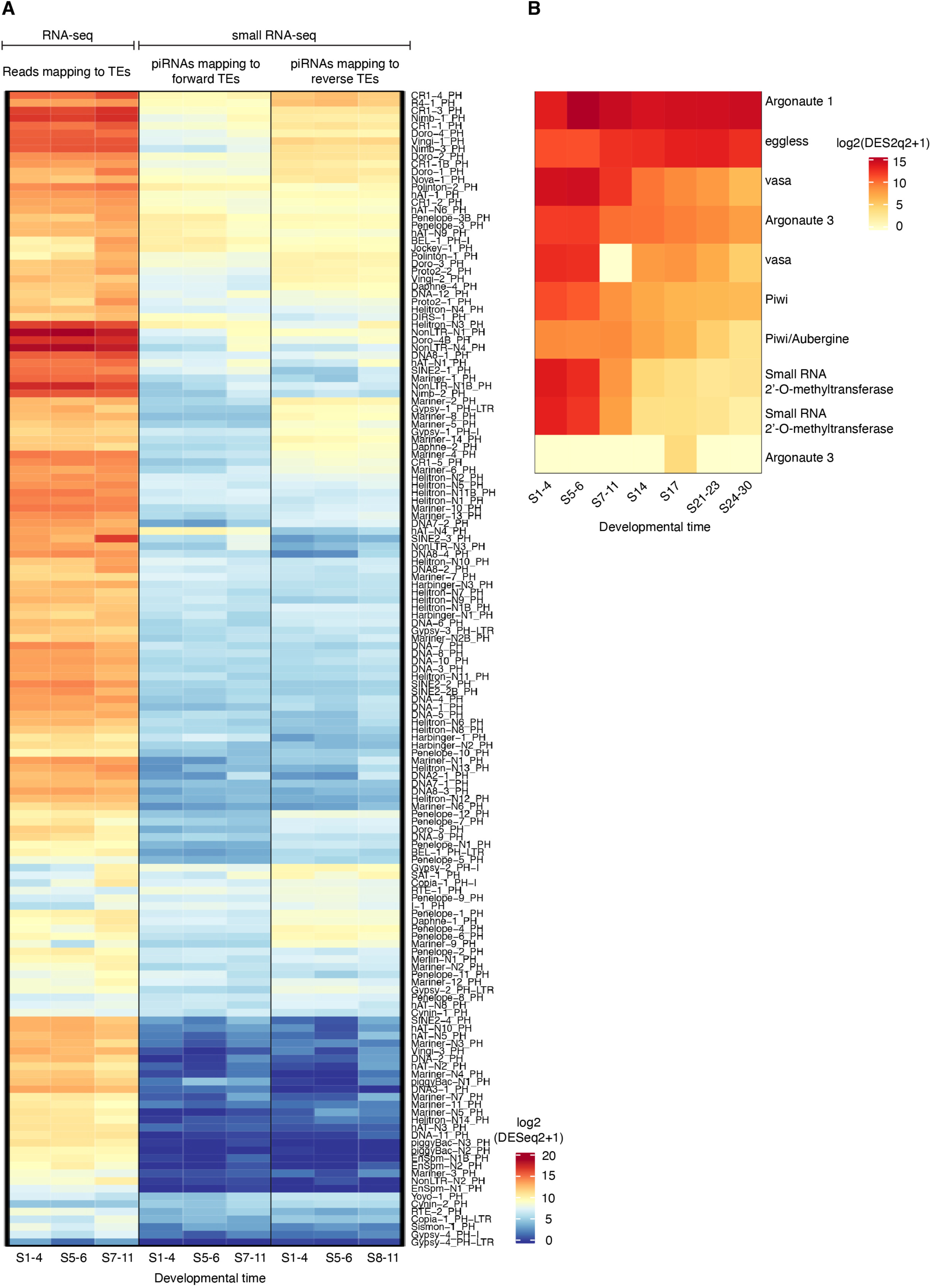
Expression analysis of transposable elements (TEs) and associated piRNAs in early embryogenesis. 151 consensus transposable elements (TEs) are listed to the right. Heatplot indicates expression level, calculated as log2 of normalized DESeq2 count+1, for each TE, and for piRNAs mapping forward and reverse to each TE, for the first three developmental time-points. TE expression was calculated from RNA-Seq data, and piRNA expression from small RNA-Seq data, from the same set of pooled embryos.

### piRNA pathway genes are highly expressed in early embryogenesis

Using our previously generated RNA-Seq data^34^ we also analysed the developmental expression profiles of mRNAs whose proteins are involved in the piRNA pathway (Figure 3B). The mRNAs of both Piwi and AGO3, whose proteins bind piRNAs and are necessary for TE silencing, are maternally loaded and highly expressed in the early embryo, consistent with a functional role at early embryonic stages. The mRNA of other proteins such as small RNA 2′O methyltransferase (required for the processing of piRNAs), and Vasa isoforms (required for the ping-pong pathway), also showed highest expression at the earliest time points. However, the mRNA of AGO1, which is involved in the siRNA and microRNA pathways but not the piRNA pathway, showed no particular enrichment at early stages, with consistently high levels throughout embryonic development. Similarly, *eggless*, which codes for a histone methyltransferase involved in heterochromatization and silencing of TEs but also other non-piRNA processes, does not show early enrichment, but consistently high expression across development.

### Expression of *piwi* and *vasa* becomes restricted to germ cells at early germband formation

Piwi and Vasa are important players for the action of primary piRNAs (Piwi-bound) and the production of piRNAs via the ping-pong cycle, respectively. If we assume conservation of the piRNA processing pathways and machinery (piwi and vasa) in *Parhyale*, it follows that cells that do not express these proteins lack the means to generate functional piRNAs. We can therefore use the temporal and spatial expression of *vasa* and *piwi* as a proxy for piRNA production in cells. A previous study of *vasa* expression during *Parhyale* development found that *vasa* mRNAs are maternally provided, while Vasa protein is only detected from the 8-cell stage onwards, and only in the germline^48,49^. To detect *piwi-* and *vasa-*positive cells, we employed a recently-developed single molecule inexpensive FISH (smiFISH) protocol^50^ that we have optimized for arthropod embryos^40^. Figure 4 shows a two-colour smiFISH stain, to detect RNAs of *piwi* and *vasa* in embryos from three different stages around germband formation (stage 8, stage 11 and stage 14). We observed that both *vasa* and *piwi* expression becomes restricted to the same small central patch of cells in the later embryo, coinciding with the time in our dataset when piRNAs are lost – between the S7-11 and S14 time-points. It therefore appears that the restriction of piRNAs to only the small set of germ cells at germband formation likely explains the apparent loss of piRNAs from the dataset at all later stages. This observation is consistent with previous findings in *Drosophila*, where piRNAs are abundant in the early embryo and contribute to processes such as the clearance of maternally provided RNAs, but are soon after only detectable in the germ cells and the surrounding somatic tissues^51–53^. Our results also align with a recent publication that failed to detect somatic piRNAs from the only crustacean surveyed to date, *A. vulgare* ^19^.

**Figure 4:**
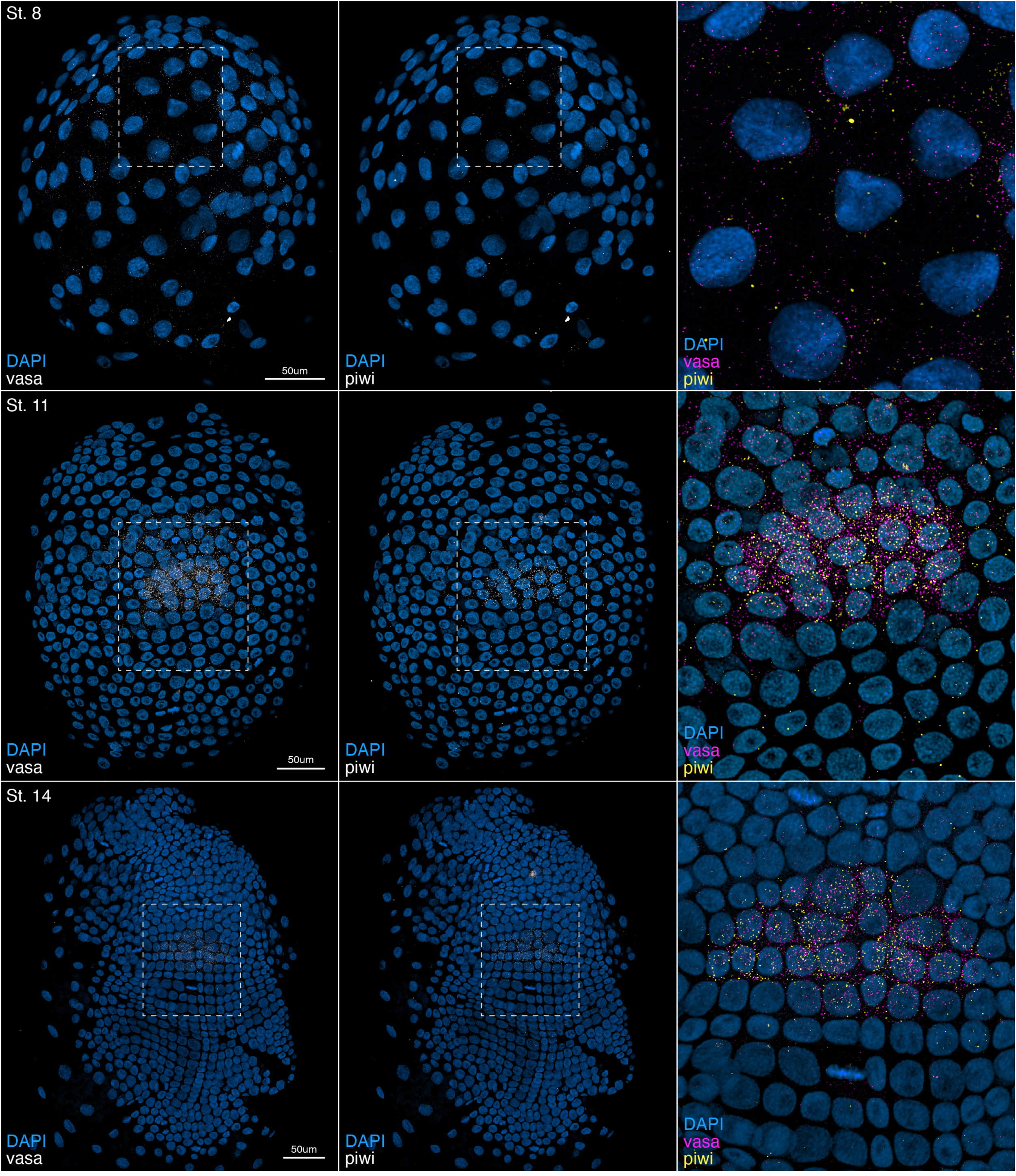
smiFISH detection of *piwi* and *vasa* RNAs in an early germband *Parhyale* embryo. Figure shows deconvoluted Z-stacked images of a stage 8, 11 and 14 of *Parhyale* embryo (anterior top) with single-molecule inexpensive fluorescent in-situ hybridization (smiFISH) staining for *piwi* and *vasa* RNAs, plus DAPI to show the nuclei. The X-flap sequence was used for both probes, labelled with CalFluor 610 (*piwi*), and Quasar 570 (*vasa*). The dashed white box indicates the region in the magnified panel.

## Discussion

In *Parhyale*, previous studies have used Vasa protein staining as a marker to identify germ cells. Vasa-positive cells are first detected during the 8-cell stage of embryogenesis (time-point S1-4 in our dataset), more precisely in the g-micromere which is known to give rise to germ cells^54^. Subsequently, four Vasa-positive cells, known as primordial germ cells (PGCs) can be detected during germ disc formation (S5-6 in our dataset), and as the disc grows these cells also undergo mitosis and increase in number. The PGCs remain clustered together through embryogenesis and by the time of hatching they have formed two clusters near the heart primordium^55^. Previous studies in *Parhyale* have also shown that the germ cells in this organism are specified by a distinct cytoplasmic region rich in germ line-associated RNAs termed the RNA-containing body (RCB)^48^. The RCB is rich in transcripts such as *vasa*, which besides its implications in piRNA biogenesis, has also been found to be necessary for germ line maintenance in *Parhyale*, but not for its formation^49^.

Our data show that the *Parhyale* embryo is pre-loaded with both TE transcripts and maternal piRNAs, the latter derived at high density from TEs and piClusters. Analysis of the expression profiles of individual TEs and their associated piRNAs across early embryogenesis found that piRNAs mapped both sense and antisense to TEs, with antisense piRNAs being more abundant. This result is consistent with piRNA-mediated silencing of TE transcripts being achieved through targeting of TEs by primary piRNAs, producing secondary piRNAs via the ping-pong pathway. Other aspects of our data also support an active ping-pong cycle in the first three time-points, spanning from the one-cell stage to germband formation. Specifically, sequence logo analysis showed a 1U/10A signature, and we found the typical 10nt 5′-5′ overlap bias indicative of ping-pong processing. These ping-pong features are strongest at the earliest time-point S1-4, which includes embryos from 1 cell to 8 cell stages. This raises the question of whether both primary and secondary piRNAs are pre-loaded into the egg, or whether only primary piRNAs are pre-loaded, and then used to produce secondary piRNAs as soon as the first copies of Vasa protein are detected, which is at the 8-cell stage^55^. In either case, it is apparent that the ping-pong mechanism is employed at the very earliest stages of embryogenesis, presumably because initial high levels of primary and secondary piRNAs are needed from the outset to defend the early embryo from the activity of TEs.

Early work in *Drosophila* showed that piRNAs are associated with the protection against TEs exclusively in the germ line. Our data show that the initially high levels of piRNAs diminish from S7-S11 onwards, and by S14, piRNAs are barely detectable regardless of their genomic origin. *vasa* and *piwi* expression serves as both a marker of the germ line cells, and a proxy for piRNA activity, since both proteins are critical for primary and secondary piRNA biogenesis. smiFISH staining of *vasa* and *piwi* RNAs in stages S8, S11 and S14 embryo highlighted that only a patch of cells, in the same zone in which germ cells have been described, were *vasa-* and *piwi*-positive, surrounded by embryonic tissues where no expression was detected.

We therefore hypothesize that in *Parhyale*, prior to PGC specification, maternally-loaded piRNAs are abundant throughout the embryo, but as the germ cells become specified and the maternal transcripts are degraded, piRNAs become restricted to just the small set of cells expressing the means for their continued production. This has been observed to be the case in *Drosophila* ^51–53^. In our bulk RNA-Seq data, the low levels of piRNAs at later embryonic stages may be explained by dilution, whereby piRNA reads originating from the few germ cells are swamped by the background of other short RNAs (e.g. microRNAs) expressed in other embryonic cells.

Together our data do not support a significant presence of somatic piRNAs in *Parhyale* beyond the earliest embryonic stages. This is consistent with findings from the only other crustacean assessed to date, *A. vulgare*. Somatic piRNAs are considered to be present in the last common ancestor of all metazoans, and also in the last common ancestor of all arthropods. Secondary losses have been reported among the arthropods, but the majority of insects studied to date display the ancestral state of somatic piRNAs. Insects evolved from crustaceans^56^, yet the two crustaceans analysed to date both lack somatic piRNAs. It is therefore likely that this loss is not shared across all crustacea, but is perhaps common to the class Malacostraca to which both *Parhyale* and *A. vulgare* belong. Future studies in additional crustacean systems are required to resolve the timing of loss of somatic piRNAs within the crustacea.

It is important to note that this study does not encompass all somatic cell types or developmental stages. piRNAs have been detected in adult insect brains (e.g. *Drosophila*), where they are thought to contribute to genome defense and neural function^18^, raising the possibility that somatic piRNA expression in *Parhyale* could occur later in development or in other tissues not sampled here.

---

**Supplementary Figure 1.**
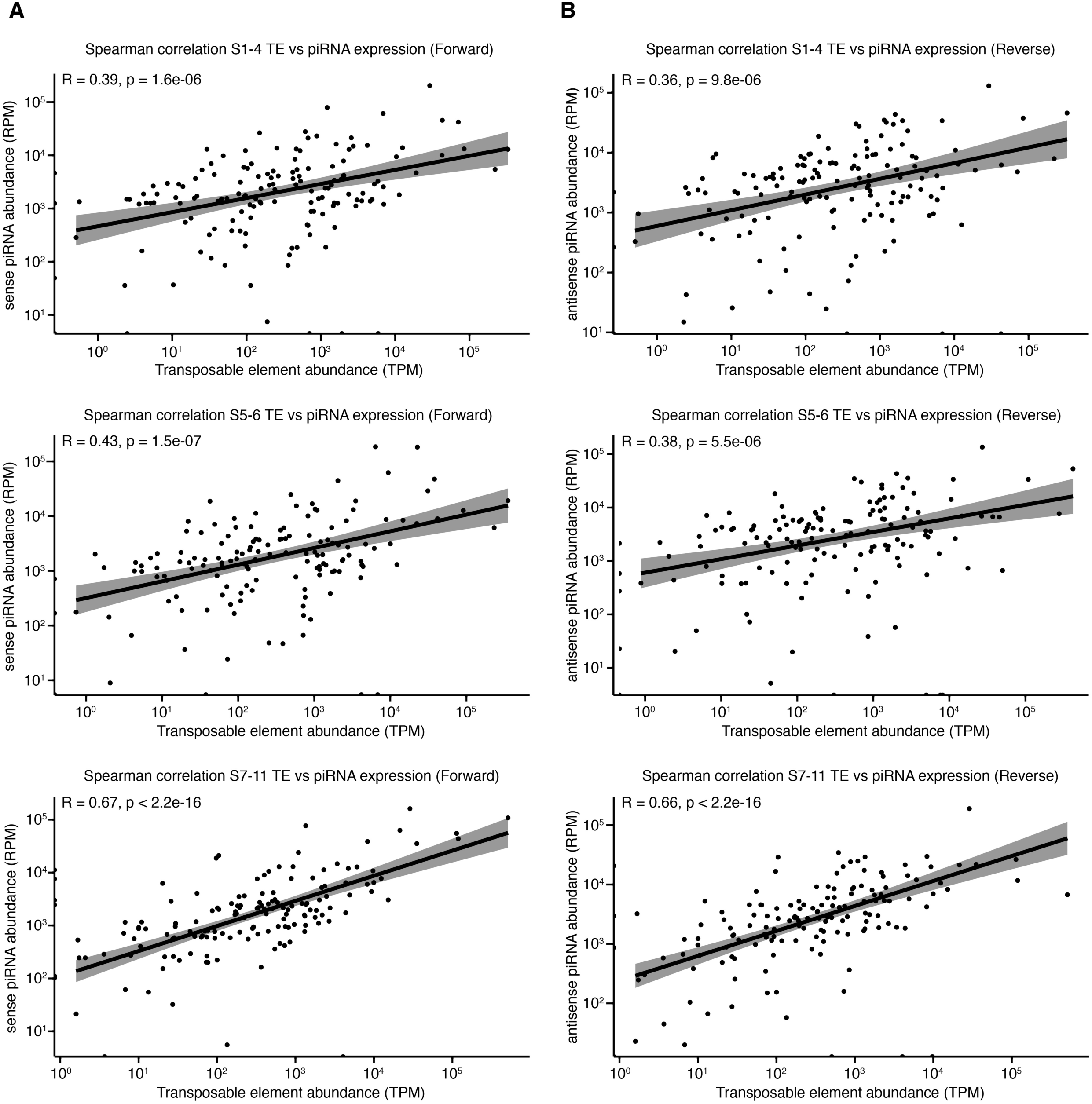
Correlation between transposable element (TE) expression and piRNA abundance in early Parhyale embryos. Scatter plots show Spearman correlations between TE transcript levels (TPM) and piRNA abundance (RPM) mapping to either the sense (A) or antisense (B) strand at stages S1–4, S5– 6, and S7–11. Correlations strengthen over time, peaking at S7–11, suggesting increased targeting of maternally loaded TEs by piRNAs as development progresses.

**Supplementary Table 1.**

**Sequences of smiFISH probes used in this study.**

List of oligonucleotide probe sequences designed for *P. hawaiensis* **piwi** and **vasa** mRNAs, used for single molecule inexpensive FISH (smiFISH) experiments.

## Acknowledgements

We thank Hilary Ashe for support and mentorship and the Wellcome Trust for a PhD studentship to L.C (203990/Z/16/A) that made this work possible.

## Author contributions

Conceptualization L.C.; culture and embryo handling L.C. and T.P.; RNA-Seq and bioinformatic analysis L.C.; in situ hybridization L.C and T.P; microscopy and image analysis T.P; writing – draft L.C. and T.P; review and editing L.C., T.P., G.Y., M.R., and S.G.J.; funding from the Wellcome Trust – PhD studentship to L.C.

## Competing interests statement

The authors declare no competing interests.

## Notes

### Competing Interest Statement

The authors have declared no competing interest.

## References

1 Aravin, A. A., Hannon, G. J. & Brennecke, J. The Piwi-piRNA pathway provides an adaptive defense in the transposon arms race. Science 318, 761–764, doi:10.1126/science.1146484 (2007).

2 Brennecke, J. et al. Discrete small RNA-generating loci as master regulators of transposon activity in Drosophila. Cell 128, 1089–1103, doi:10.1016/j.cell.2007.01.043 (2007).

3 Gunawardane, L. S. et al. A slicer-mediated mechanism for repeat-associated siRNA 5’ end formation in Drosophila. Science 315, 1587–1590, doi:10.1126/science.1140494 (2007).

4 Bergman, C. M., Quesneville, H., Anxolabehere, D. & Ashburner, M. Recurrent insertion and duplication generate networks of transposable element sequences in the Drosophila melanogaster genome. Genome Biol 7, R112, doi:10.1186/gb-2006-7-11-r112 (2006).

5 Malone, C. D. & Hannon, G. J. Small RNAs as guardians of the genome. Cell 136, 656–668, doi:10.1016/j.cell.2009.01.045 (2009).

6 Horwich, M. D. et al. The Drosophila RNA methyltransferase, DmHen1, modifies germline piRNAs and single-stranded siRNAs in RISC. Curr Biol 17, 1265–1272, doi:10.1016/j.cub.2007.06.030 (2007).

7 Saito, K. et al. Pimet, the Drosophila homolog of HEN1, mediates 2’-O-methylation of Piwi-interacting RNAs at their 3’ ends. Genes Dev 21, 1603–1608, doi:10.1101/gad.1563607 (2007).

8 Brower-Toland, B. et al. Drosophila PIWI associates with chromatin and interacts directly with HP1a. Genes Dev 21, 2300–2311, doi:10.1101/gad.1564307 (2007).

9 Rangan, P. et al. piRNA production requires heterochromatin formation in Drosophila. Curr Biol 21, 1373–1379, doi:10.1016/j.cub.2011.06.057 (2011).

10 Le Thomas, A. et al. Piwi induces piRNA-guided transcriptional silencing and establishment of a repressive chromatin state. Genes Dev 27, 390–399, doi:10.1101/gad.209841.112 (2013).

11 Lim, A. K. & Kai, T. Unique germ-line organelle, nuage, functions to repress selfish genetic elements in Drosophila melanogaster. Proc Natl Acad Sci U S A 104, 6714–6719, doi:10.1073/pnas.0701920104 (2007).

12 Xiol, J. et al. RNA clamping by Vasa assembles a piRNA amplifier complex on transposon transcripts. Cell 157, 1698–1711, doi:10.1016/j.cell.2014.05.018 (2014).

13 Robine, N. et al. A broadly conserved pathway generates 3’UTR-directed primary piRNAs. Curr Biol 19, 2066–2076, doi:10.1016/j.cub.2009.11.064 (2009).

14 Barckmann, B. et al. Aubergine iCLIP Reveals piRNA-Dependent Decay of mRNAs Involved in Germ Cell Development in the Early Embryo. Cell Rep 12, 1205–1216, doi:10.1016/j.celrep.2015.07.030 (2015).

15 Rouget, C. et al. Maternal mRNA deadenylation and decay by the piRNA pathway in the early Drosophila embryo. Nature 467, 1128–1132, doi:10.1038/nature09465 (2010).

16 Toth, K. F., Pezic, D., Stuwe, E. & Webster, A. The piRNA Pathway Guards the Germline Genome Against Transposable Elements. Adv Exp Med Biol 886, 51–77, doi:10.1007/978-94-017-7417-8_4 (2016).

17 Malone, C. D. et al. Specialized piRNA pathways act in germline and somatic tissues of the Drosophila ovary. Cell 137, 522–535, doi:10.1016/j.cell.2009.03.040 (2009).

18 Perrat, P. N. et al. Transposition-driven genomic heterogeneity in the Drosophila brain. Science 340, 91–95, doi:10.1126/science.1231965 (2013).

19 Lewis, S. H. et al. Pan-arthropod analysis reveals somatic piRNAs as an ancestral defence against transposable elements. Nat Ecol Evol 2, 174–181, doi:10.1038/s41559-017-0403-4 (2018).

20 Wang, W., Ashby, R., Ying, H., Maleszka, R. & Foret, S. Contrasting Sex-and Caste-Dependent piRNA Profiles in the Transposon Depleted Haplodiploid Honeybee Apis mellifera. Genome Biol Evol 9, 1341–1356, doi:10.1093/gbe/evx087 (2017).

21 Kiuchi, T. et al. A single female-specific piRNA is the primary determiner of sex in the silkworm. Nature 509, 633–636, doi:10.1038/nature13315 (2014).

22 Ninova, M., Griffiths-Jones, S. & Ronshaugen, M. Abundant expression of somatic transposon-derived piRNAs throughout Tribolium castaneum embryogenesis. Genome Biol 18, 184, doi:10.1186/s13059-017-1304-1 (2017).

23 Gainetdinov, I., Colpan, C., Arif, A., Cecchini, K. & Zamore, P. D. A Single Mechanism of Biogenesis, Initiated and Directed by PIWI Proteins, Explains piRNA Production in Most Animals. Mol Cell 71, 775–790 e775, doi:10.1016/j.molcel.2018.08.007 (2018).

24 Llonga, N., Ylla, G., Bau, J., Belles, X. & Piulachs, M. D. Diversity of piRNA expression patterns during the ontogeny of the German cockroach. J Exp Zool B Mol Dev Evol 330, 288–295, doi:10.1002/jez.b.22815 (2018).

25 Mondal, M., Mansfield, K. & Flynt, A. siRNAs and piRNAs collaborate for transposon control in the two-spotted spider mite. RNA 24, 899–907, doi:10.1261/rna.065839.118 (2018).

26 La Greca, A. et al. PIWI-interacting RNAs are differentially expressed during cardiac differentiation of human pluripotent stem cells. PLoS One 15, e0232715, doi:10.1371/journal.pone.0232715 (2020).

27 Vogt, G. in Encyclopedia of Reproductio Vol. 6 (2018).

28 Bar-On, Y. M., Phillips, R. & Milo, R. The biomass distribution on Earth. Proc Natl Acad Sci U S A 115, 6506–6511, doi:10.1073/pnas.1711842115 (2018).

29 Martin, M. Cutadapt removes adapter sequences from high-throughput sequencing reads. EMBnet.journal 17, 10–12, doi:10.14806/ej.17.1.200 (2011).

30 Langmead, B., Trapnell, C., Pop, M. & Salzberg, S. L. Ultrafast and memory-efficient alignment of short DNA sequences to the human genome. Genome Biology 10, R25, doi:10.1186/gb-2009-10-3-r25 (2009).

31 Consortium, R. N. RNAcentral 2021: secondary structure integration, improved sequence search and new member databases. Nucleic Acids Res 49, D212–D220, doi:10.1093/nar/gkaa921 (2021).

32 Calvo, L., Birgaoanu, M., Pettini, T., Ronshaugen, M. & Griffiths-Jones, S. The developmental transcriptome of <em>Parhyale hawaiensis</em>: microRNAs and mRNAs show different expression dynamics during the maternal-zygotic transition. bioRxiv, 2021.2006.2025.449901, doi:10.1101/2021.06.25.449901 (2021).

33 Rosenkranz, D. & Zischler, H. proTRAC--a software for probabilistic piRNA cluster detection, visualization and analysis. BMC Bioinformatics 13, 5, doi:10.1186/1471-2105-13-5 (2012).

34 Calvo, L., Birgaoanu, M., Pettini, T., Ronshaugen, M. & Griffiths-Jones, S. The embryonic transcriptome of Parhyale hawaiensis reveals different dynamics of microRNAs and mRNAs during the maternal-zygotic transition. Sci Rep 12, 174, doi:10.1038/s41598-021-03642-9 (2022).

35 Antoniewski, C. Computing siRNA and piRNA overlap signatures. Methods Mol Biol 1173, 135–146, doi:10.1007/978-1-4939-0931-5_12 (2014).

36 Liao, Y., Smyth, G. K. & Shi, W. featureCounts: an efficient general purpose program for assigning sequence reads to genomic features. Bioinformatics 30, 923–930, doi:10.1093/bioinformatics/btt656 (2014).

37 Love, M. I., Huber, W. & Anders, S. Moderated estimation of fold change and dispersion for RNA-seq data with DESeq2. Genome Biol 15, 550, doi:10.1186/s13059-014-0550-8 (2014).

38 Wickham, H. ggplot2: Elegant Graphics for Data Analysis. (2016).

39 Gu, Z., Eils, R. & Schlesner, M. Complex heatmaps reveal patterns and correlations in multidimensional genomic data. Bioinformatics 32, 2847–2849, doi:10.1093/bioinformatics/btw313 (2016).

40 Calvo, L., Ronshaugen, M. & Pettini, T. smiFISH and embryo segmentation for single-cell multi-gene RNA quantification in arthropods. Commun Biol 4, 352, doi:10.1038/s42003-021-01803-0 (2021).

41 Vagin, V. V. et al. A distinct small RNA pathway silences selfish genetic elements in the germline. Science 313, 320–324, doi:10.1126/science.1129333 (2006).

42 Aravin, A. et al. A novel class of small RNAs bind to MILI protein in mouse testes. Nature 442, 203–207, doi:10.1038/nature04916 (2006).

43 Watanabe, T. & Lin, H. Posttranscriptional regulation of gene expression by Piwi proteins and piRNAs. Mol Cell 56, 18–27, doi:10.1016/j.molcel.2014.09.012 (2014).

44 Andersen, P. R., Tirian, L., Vunjak, M. & Brennecke, J. A heterochromatin-dependent transcription machinery drives piRNA expression. Nature 549, 54–59, doi:10.1038/nature23482 (2017).

45 Yamanaka, S., Siomi, M. C. & Siomi, H. piRNA clusters and open chromatin structure. Mob DNA 5, 22, doi:10.1186/1759-8753-5-22 (2014).

46 Sun, Y. H. et al. Domestic chickens activate a piRNA defense against avian leukosis virus. Elife 6, doi:10.7554/eLife.24695 (2017).

47 Weick, E. M. & Miska, E. A. piRNAs: from biogenesis to function. Development 141, 3458–3471, doi:10.1242/dev.094037 (2014).

48 Gupta, T. & Extavour, C. G. Identification of a putative germ plasm in the amphipod Parhyale hawaiensis. Evodevo 4, 34, doi:10.1186/2041-9139-4-34 (2013).

49 Ozhan-Kizil, G., Havemann, J. & Gerberding, M. Germ cells in the crustacean Parhyale hawaiensis depend on Vasa protein for their maintenance but not for their formation. Dev Biol 327, 230–239, doi:10.1016/j.ydbio.2008.10.028 (2009).

50 Tsanov, N. et al. smiFISH and FISH-quant - a flexible single RNA detection approach with super-resolution capability. Nucleic Acids Res 44, e165, doi:10.1093/nar/gkw784 (2016).

51 Renault, A. D. vasa is expressed in somatic cells of the embryonic gonad in a sex-specific manner in Drosophila melanogaster. Biol Open 1, 1043–1048, doi:10.1242/bio.20121909 (2012).

52 Shigenobu, S., Kitadate, Y., Noda, C. & Kobayashi, S. Molecular characterization of embryonic gonads by gene expression profiling in Drosophila melanogaster. Proc Natl Acad Sci U S A 103, 13728–13733, doi:10.1073/pnas.0603767103 (2006).

53 Marie, P. P., Ronsseray, S. & Boivin, A. From Embryo to Adult: piRNA-Mediated Silencing throughout Germline Development in Drosophila. G3 (Bethesda) 7, 505–516, doi:10.1534/g3.116.037291 (2017).

54 Gerberding, M., Browne, W. E. & Patel, N. H. Cell lineage analysis of the amphipod crustacean Parhyale hawaiensis reveals an early restriction of cell fates. Development 129, 5789–5801, doi:10.1242/dev.00155 (2002).

55 Extavour, C. G. The fate of isolated blastomeres with respect to germ cell formation in the amphipod crustacean Parhyale hawaiensis. Dev Biol 277, 387–402, doi:10.1016/j.ydbio.2004.09.030 (2005).

56 Lozano-Fernandez, J., et al. Pancrustacean Evolution Illuminated by Taxon-Rich Genomic-Scale Data Sets with an Expanded Remipede Sampling. Genome Biol Evol 11, 2055–2070, doi:10.1093/gbe/evz097 (2019).

